# Stick-slip motion and universal statistics of cargo transport within living cells

**DOI:** 10.1101/2025.05.19.654995

**Authors:** Yusheng Shen, Caishan Yan, Pingbo Huang, Kassandra M. Ori-McKenney, Pik-Yin Lai, Penger Tong

**Affiliations:** Department of Physics, Hong Kong University of Science and Technology, Clear Water Bay, Kowloon, Hong Kong; Division of Life Science and State Key Laboratory of Molecular Neuroscience, Hong Kong University of Science and Technology, Clear Water Bay, Kowloon, Hong Kong; Department of Molecular and Cellular Biology, University of California, Davis, Davis, CA 95616, USA; Department of Physics and Center for Complex Systems, National Central University, Taoyuan City 320, Taiwan; Physics Division, National Center for Theoretical Sciences, Taipei 10617, Taiwan

## Abstract

Cargo transport within cells is a vital biological process that relies on the intricate interplay between motor proteins, microtubules, and the complex intracellular environment. In this study, we unveil a universal transport mechanism characterized by stick-slip motion, which governs the dynamics of intracellular vesicle transport. By analyzing a comprehensive dataset of vesicle trajectories across various cell types and intracellular environments, we demonstrate that the cargo velocities consistently follow a Gamma distribution, revealing a common statistical pattern amidst the diversity of biological cargoes. Our experimental findings are well-described by a theoretical model that connects the Brownian-correlated kinetic friction between motor-cargo complexes and their surroundings to the observed universal Gamma distribution of cargo velocities. This model elucidates the stick-slip dynamics governing intracellular cargo transport, which are pertinent to various cellular processes such as vesicle budding, organelle transport, and cell migration.

---

Our understanding of molecular motions began with simple gas molecules at equilibrium, which possess kinetic energy and have velocities that follow the Maxwell-Boltzmann distribution, a Gaussian function in any given direction [1]. In contrast, complex molecules or molecular assemblies, such as motor proteins like myosin, kinesin, and dynein, can convert chemical energy from adenosine triphosphate (ATP) hydrolysis into directed motion [2]. As these motor proteins continuously consume chemical energy during movement, they exist in a non-equilibrium state. In in vitro systems, purified motor proteins exhibit unrestricted directed motion along actin filaments or microtubules, and their velocities display a narrow distribution that is accurately described by a Gaussian function [3–9]. Gaussian distributions have also been observed in vitro for groups of motors transporting cargoes (or vesicles) [10–13].

In living cells, motor proteins cluster together to transport cargo along the cytoskeletal network, facilitating the distribution of cellular components over long distances. These motor-driven vesicles must navigate a complex intracellular environment that is crowded, viscoelastic, sticky, and heterogeneous. The interior of a cell is filled with macromolecules and entangled networks of organelles and cytoskeletal polymers, which can occupy 10-40% of the total cellular volume [14–16]. These components form a porous and viscoelastic medium that hinders the movement of large particles, such as vesicles and mitochondria [17–19]. Moreover, the motor-cargo complexes involved in trafficking are not inert; they interact extensively with the local cytoskeleton and organelles, including actin-rich regions and the endoplasmic reticulum (ER) [20–25].

As a result, intracellular cargo transport often exhibits random changes in direction, alternating between directed “runs” and diffusive jiggling movements or “pauses” [21, 22, 26–31]. These pauses can account for up to 80% of the total travel time. The cargoes include secretory vesicles, synaptic vesicles, signaling endosomes, mitochondria, and lipid droplets, all of which vary in shape and size. The cargo velocities have been observed to deviate from a Gaussian distribution [10, 28, 30, 32–35], but the precise functional form of the velocity distribution remains undetermined due to insufficient statistical sampling. It remains an intriguing question whether the moving cargoes in a cell follow a universal velocity distribution, and if so, what the underlying physical mechanisms are.

In this study, we systematically investigate cargo transport across various vesicle types, cell types, and intracellular environments. Utilizing advanced single-molecule tracking and sorting algorithms, we collect an extensive dataset of vesicle trajectories from over 450 live cells, covering a wide range of sampling rates and distances across the entire cell. Our statistical analysis reveals that, unlike in vitro systems where cargo velocities follow Gaussian statistics, the velocities of cargo in living cells consistently adhere to a Gamma distribution across diverse conditions. This robust statistical pattern highlights a universal transport mechanism applicable to a variety of vesicles. Our theoretical model provides a quantitative coarse-grained description of the stick-slip motion of motor-cargo complexes in cells. This model envisions that as a cargo moves through the porous, viscoelastic cytoplasm, the motor pulling force must overcome both the viscous drag from the surrounding mobile protein solution and the elastic friction resulting from interactions with the surrounding cytoskeletal and membrane networks. The continuous pinning and depinning (or stick-slip) of the interaction bonds between the motor-cargo complex and the surrounding protein networks lead to a universal Gamma distribution of velocities.

## Results

We first investigate the endocytic trafficking of epidermal growth factor receptors (EGFR) in BEAS-2B cells, a commonly used human lung epithelial cell line [36]. The EGFRs on the cell membrane are labeled with bright, photostable EGF-conjugated quantum dots (EGF-QDs) at room temperature. When activated by EGF-QDs at 37^°^C, the EGFR-EGF-QD complexes form endocytic vesicles that travel through the endosomal system, progressing from early to late endosomes before being degraded in lysosomes [37]. The motor-driven EGFR-containing endosomes, hereafter referred to as EGFR-endosomes, serve as a well-characterized model for studying cargo transport. Using a homemade single-particle tracking (SPT) program with approximately 20 nm spatial resolution [38, 39], we track the position **r**(*t*) of the EGFR-endosomes over long time *t*. This algorithm enables us to collect over 100,000 individual vesicle trajectories from more than 454 cells, capturing their movements across the entire cell (see Table S1 and Supplementary Information (SI) Sec. I. for more details [40]).

Figure 1(a) shows a collection of mobile trajectories of the late-stage EGFR-endosomes (more than 40 minutes after activation by EGF-QDs). These EGFR-endosomes display considerable dynamic heterogeneity, as indicated by the significant variations in trajectory shapes, sizes and their position-dependence relative to the cell nucleus. Some endosomes move slowly with a short trajectory primarily in the perinuclear region, while others travel quickly with longer paths that span the entire cell. Overall, the trajectories exhibit a radial pattern around the nucleus, resembling the microtubule network in BEAS-2B cells. As illustrated in Figs. 1(b) and 1(c), individual vesicle trajectories show a complex motility pattern, characterized by intermittent runs (“on-state”), pauses (“off-state”), and reversals. The “on-state” refers to directed motion of the EGFR-endosome along relatively straight and long paths, while the “off-state” describes jiggling or diffusion-like motion with minimal displacement. This “run-and-tumble” movement has also been observed in various motor-driven cargoes, including protein-specific endosomes, lysosomes, secretory vesicles, and artificial beads [21, 22, 28–30]. These findings suggest that the two-state motion of motor-driven vesicles in cells may stem from a common underlying mechanism.

**FIG. 1.**
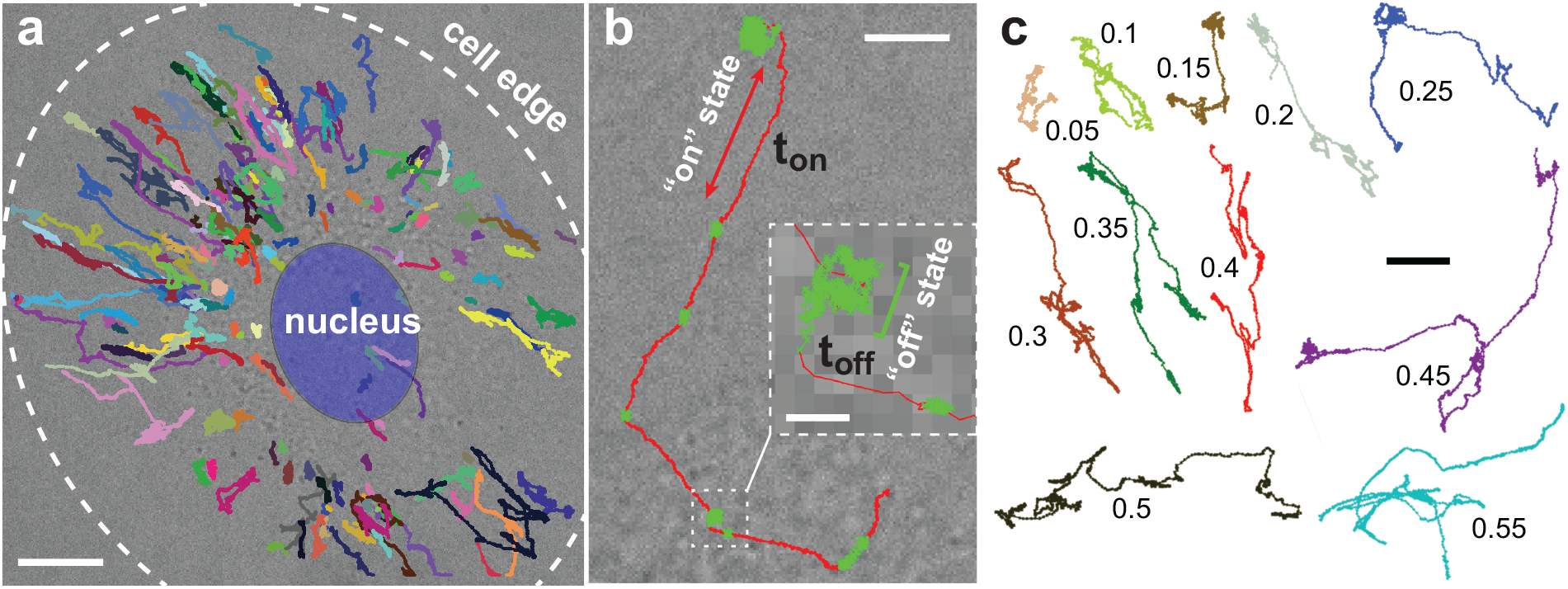
Cargo transport across a living cell. (a) 194 representative mobile trajectories (colored lines) superimposed on a simultaneously taken bright-field image of a BEAS-2B cell, illustrating the intracellular motion of the late-stage EGFR-endosomes over 1800 time steps (180 s). The white dashed circle marks the cell periphery, while the blue ellipse indicates the nucleus. The scale bar is 10 *μ*m. (b) A representative trajectory of an EGFR-endosome, highlighting intermittent switching between the on-state (red) and off-state (green). The inset provides a magnified view of a section of the trajectory. The scale bars: 5 *μ*m (main image) and 0.5 *μ*m (inset). (c) Eleven representative mobile trajectories of EGFR-endosomes with varying on-state probabilities *p*on, as marked by a nearby number. The scale bar: 5 *μ*m.

To analyze the statistical properties of the on-state and off-state trajectory segments separately, we employ an automated selection algorithm [41] to distinguish the two types of segments within each trajectory (see SI Fig. S1 and Sec. I for details [40]). Once identified, we label each vesicle trajectory by its on-state probability,

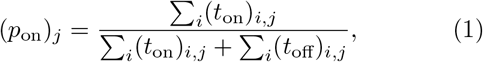

where (*t*_on_)_*i*,*j*_ and (*t*_off_)_*i*,*j*_ are the dwell times of the *i*th on-state and off-state segments, respectively; both belong to the *j*th trajectory. For simplicity, we will drop the subscript *j*. This allows us to categorize the vesicle trajectories into two groups: an immobile group with *p*_on_ = 0 and a mobile group with *p*_on_ *>* 0, which comprises ~42% of the trajectories. We focus on the mobile trajectories that exhibit clear motor-driven directed motion. Figure 1(c) displays eleven representative mobile trajectories with varying values of *p*_on_, illustrating the characteristic shapes of EGFR-endosome trajectories. The *p*_on_ values range from 0 to 0.85, with a mean of ⟨*p*_on_⟩ ≃ 0.12 for mobile trajectories from different BEAS-2B cells, consistent with previous findings [21].

From the large volume of the extracted on-state segments, we analyze the statistics of three on-state velocities at different time scales. The instantaneous velocity is defined as *v*_ins_ = Δ*r/*Δ*t*, where Δ*r* is the distance traveled by the vesicle over the shortest sample time (time stamp), Δ*t* = 0.1 s. The segment velocity *v*_seg_ is the average of *v*_ins_ over each segment, which typically lasts between 1.3 and 4.2 s. The trajectory velocity *v*_tra_ is the average of *v*_seg_ across multiple segments within each trajectory. Figure 2(a) shows the obtained probability density functions (PDFs) of *v*_tra_ and *v*_seg_ for EGFR-endosomes in BEAS-2B cells. The measured *P*(*v*_tra_) shows a narrower distribution compared to *P*(*v*_seg_), as fluctuations in *v*_seg_ are partially averaged out in *v*_tra_. The obtained values of *v*_*tra*_ are consistent with previous reports, ranging from 0.88 to 2 *μ*m/s for EGFR-endosomes and lysosomes/late endosomes [42, 43].

**FIG. 2.**
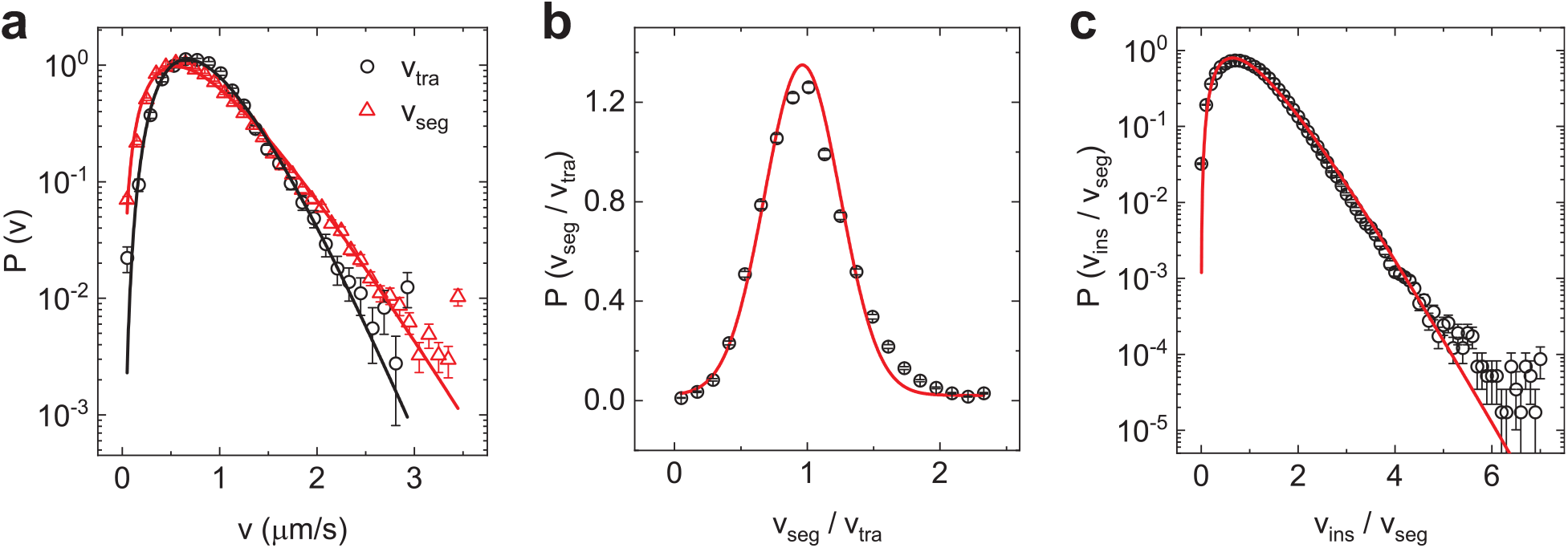
Statistics of on-state velocities. (a) Measured probability density functions (PDFs) of trajectory velocity *v*_tra_ (black circles) and segment velocity *v*_seg_ (red triangles) for EGFR-endosomes in BEAS-2B cells. The black and red solid lines show the best fits of Eq. (2) to the data, with fitting parameters listed in Table S2. (b) Measured PDF *P*(*v*_seg_/*v*_tra_) of the normalized segment velocity, *v*_seg_/*v*_tra_, for EGFR-endosomes. The red solid line shows a Gaussian fit, *P*(*v*) *∝*exp[− (*v* − *μ*)^2^/(2*σ*^2^)], with mean value *μ* and standard deviation *σ* provided in Table S2. (c) Measured PDF *P*(*v*_ins_/*v*_seg_) of the normalized instantaneous velocity *v*_ins_/*v*_seg_ for EGFR-endosomes. The red solid line shows the best fit of Eq. (2) to the data, with fitting parameters given in Table S2.

To further understand the differences between *P*(*v*_tra_) and *P*(*v*_seg_), we replot *P*(*v*_seg_) in Fig. 2(b) as a function of the normalized segment velocity, *v*_seg_/*v*_tra_ *≡ v*_seg_/(*v*_seg_)_*i*_, where ⟨*v*_seg_⟩ _*i*_ is the mean velocity of the *i*th trajectory. This normalization removes velocity fluctuations among different trajectories, allowing us to focus on variations among segments within the same trajectory. Figure 2(b) shows that the PDF *P*(*v*_seg_/*v*_tra_) centers around unity with a narrow peak, well-described by a Gaussian distribution (red solid line) with a mean *μ ≃* 1 and a small standard deviation *σ* = 0.64. Thus, Fig. 2(a) and 2(b) suggest that the velocity variations in Fig. 2(a) primarily arise from differences between trajectories (or motor-cargo assemblies), while variations among different segments within the same trajectory (or assembly) are relatively small, despite intermittent switching between the on-state and off-state. In other words, the motor-cargo assembly linked to each trajectory remains stable while transporting over long distances through the varied cellular environment.

Similarly, Fig. 2(c) shows the measured PDF of the normalized instantaneous velocity *v*_ins_/*v*_seg_. The PDF *P*(*v*_ins_/*v*_seg_) shows a broad distribution, peaking at *v*_ins_/*v*_seg_ = 0.67 and followed by a long exponential-like tail that lasts up to 9 times the peak value. Figure 2 thus demonstrates that, unlike in-vitro motor proteins, motor-driven EGFR-endosomes in living cells do not undergo smooth, unrestricted motion in the on-state. Their velocities exhibit substantial variations, both spatially among different EGFR-endosomes (i.e., *v*_tra_ for different trajectories) and temporally for the same EGFR-endosome (i.e., *v*_ins_ at different times).

The two sets of data in Fig. 2(a) both exhibit an asymmetrical distribution with a long exponential-like tail, which can be effectively modeled by the Gamma distribution [44],

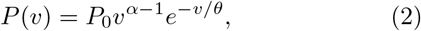

where *α* and *θ* are two fitting parameters, and *P*_0_ = 1*/*(*θ*^*α*^Γ(*α*)) is a normalization factor with Γ(*α*) being the Gamma function (see SI Sec. III.A. for more details [40]). As shown in Fig. 2(a), the Gamma distribution (solid lines) fits the data well, with the fitting parameters listed in Table S2. Figure 2(c) indicates that the PDF *P*(*v*_ins_*/v*_seg_) also follows the Gamma distribution (solid line). Thus, Fig. 2 demonstrates that velocity fluctuations among different EGFR-endosomes, as well as for the same endosome over time, all adhere to the Gamma distribution.

We now examine whether the Gamma distribution applies to other motor-driven vesicles, including lysosomes, early endosomes, and secretory vesicles. These vesicles interact differently with cellular networks and are transported by distinct sets of motor proteins [20, 30, 45, 46]. Their motion in BEAS-2B cells is recorded and analyzed in a manner similar to that described above (see SI Sec. I for more details [40]). Figure 3(a) shows the measured PDFs *P*(*v*_tra_) for EGFR-endosomes (black circles, re-plotted for comparison), LAMP1-positive lysosomes (red triangles), Rab5-positive early endosomes (blue diamonds), and Rab6A-positive secretory vesicles (green pentagons). Despite their varied interactions with the local environment and different biochemical origins and functions, the PDFs *P*(*v*_tra_) for all four vesicle types are well described by the Gamma distribution (solid lines). The fitting parameters are provided in Table S2.

**FIG. 3.**
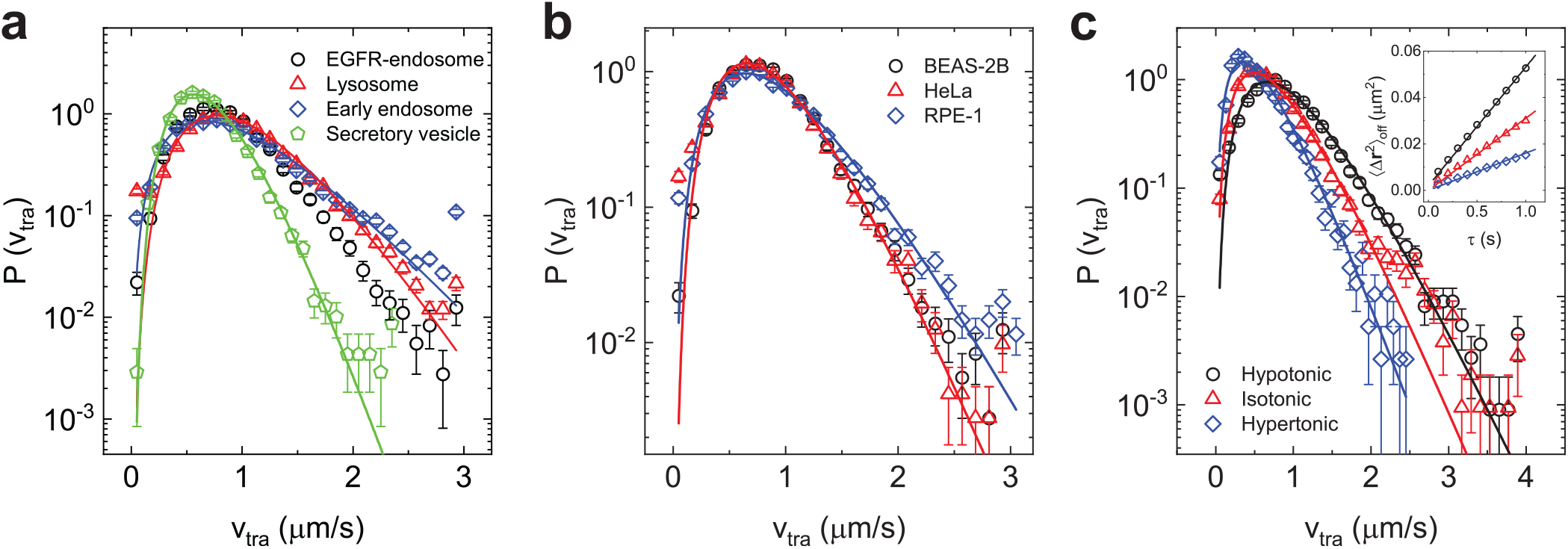
Universal statistics of cargo transport for various vesicles, cell types, and intracellular crowdednesses. (a) Measured PDFs *P*(*v*_tra_) of trajectory velocity *v*_tra_ for EGFR-endosomes (black circles, re-plot from Fig. 2(a) for comparison), lysosomes (red triangles), early endosomes (blue diamonds), and secretory vesicles (green pentagons) in BEAS-2B cells. The solid colored lines show the best fits of Eq. (2) to the data, with fitting parameters listed in Table S2. (b) Measured PDFs *P*(*v*_tra_) for EGFR-endosomes in three cell types: BEAS-2B cells (black circles), HeLa cells (red triangles), and RPE-1 cells (blue diamonds), with corresponding fits shown. (c) Measured PDFs *P*(*v*_tra_) for EGFR-endosomes in BEAS-2B cells treated with hypotonic (250 mOsm, black circles), isotonic (310 mOsm, red triangles), and hypertonic (400 mOsm, blue diamonds) solutions, along with fits. The inset shows the measured mean-squared displacement (MSD), ⟨Δ*r*^2^(*τ*)⟩ _off_, as a function of delay time *τ* for EGFR-endosomes in the off-state, with BEAS-2B cells treated with the different solutions. The solid lines show linear fits, 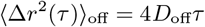, where *D*_off_ is the effective diffusion coefficient in the off-state.

We also investigate how the velocity statistics of EGFR-endosomes vary with their host cells (see SI Sec. I for details [40]). Figure 3(b) shows the measured PDFs *P*(*v*_tra_) for EGFR-endosomes in BEAS-2B cells (black circles), human cervical cancer cell line (HeLa cells, red triangles) and human retinal pigment epithelial cells (RPE-1, blue diamonds). The measured PDFs *P*(*v*_tra_) for EGFR-endosomes in three cell types are all well described by the Gamma distribution (solid lines). Comparing Fig. 3(b) with Fig. 3(a), we observe that the PDFs *P*(*v*_tra_) for different vesicles within the same cell type (BEAS-2B) show significantly greater variability than those for the same vesicle (EGFR-endosomes) but in different cell types, which exhibit minimal changes.

Furthermore, we explore how the velocity statistics of EGFR-endosomes change when varying cell volume by exposing BEAS-2B cells to culture media of different osmolarities. In hypotonic solutions, cells swell due to water intake, while in hypertonic solutions, they shrink from water loss. Figure 3(c) shows the measured PDFs *P*(*v*_tra_) for EGFR-endosomes in BEAS-2B cells treated with hypotonic, isotonic, and hypertonic solutions (see SI Sec. I for details [40]). All three datasets exhibit similar shape and are well described by the Gamma distribution (solid lines). As cell volume increases, the most probable (peak) value of *P*(*v*_tra_) also rises. When the velocity variable is normalized by its mean value ⟨*v*_tra_⟩ obtained under different osmotic conditions, the resulting PDFs *P*(*v*_tra_*/* ⟨*v*_tra_⟩) collapse onto a common Gamma distribution curve (see Fig. S3(c) in SI [40]). Thus, Fig. 3 demonstrates that the observed Gamma distribution of trajectory velocity *v*_tra_ is a robust feature of cargo transport, invariant across different vesicles, cell types, and level of intracellular crowding.

While changes in cell volume may lead to some biochemical alterations [31, 47], their primary effect is on the mechanical properties of the cytoplasm through which motor-driven vesicles travel. As cell volume decreases, the cytoplasm becomes more crowded, increasing its effective viscosity and viscoelasticity. The inset in Fig. 3(c) shows the measured mean squared displacements (MSDs), ⟨Δ*r*^2^(*τ*) ⟩ _off_, of EGFR-endosomes in the off-state for BEAS-2B cells treated with the hypotonic, isotonic, or hypertonic solutions. Here, the two-dimensional displacement vector is defined as, Δ**r**(*τ*) = **r**(*t* + *τ*) − **r**(*t*), where *τ* is the delay time. All MSD curves are well described by the linear function, (Δ*r*^2^(*τ*))_off_ = 4*D*_off_*τ*, indicating that EGFR-endosomes in the off-state undergo diffusion-like motion. The obtained effective diffusion coefficient *D*_off_ decreases from 0.014 to 0.0077 and then to 0.0040 *μm*^2^/*s*, as cell volume decreases.

In addition to analyzing trajectory velocity statistics, we compute the PDFs *P*(*v*_seg_/*v*_tra_) and *P*(*v*_ins_*/v*_seg_) for various vesicles in BEAS-2B cells and for EGFR-endosomes in three cell types. As shown in Fig. S3(a) in SI [40], the obtained PDFs *P*(*v*_seg_/*v*_tra_) all collapse into a common Gaussian distribution. Similarly, the obtained PDFs *P*(*v*_ins_/*v*_seg_) converge to a common Gamma distribution, as shown in Fig. S3(b) in SI [40]. These findings are consistent with those presented in Fig. 2 (see SI Sec. II.A. for more details [40]).

## Discussion

As the cargo-motor complex navigates through the cytoplasm, it encounters resistance from both the viscous effects of the surrounding mobile protein solution and the elastic effects of the protein networks. The viscous resistance is described by a drag force *γ*(*dx/dt*), where *γ* is the drag coefficient that depends on the size of the cargo and the viscosity of the cytoplasm, while *v*(*t*) = *dx/dt* represents the speed of the cargo. In contrast, the elastic resistance is modeled as a pinning force field *F*_pin_(*x*), which arises from interactions with the surrounding protein networks. Examples of these interactions include (i) physical tethering of vesicles to the endoplasmic reticulum (ER) through membrane contact sites [20, 48]; (ii) physical obstructions caused by actin patches and microtubule intersections [21, 22, 25]; and (iii) opposing forces exerted by different motors on the same cargo (referred to as a “tug of war”) or by motors interacting with microtubules oriented in opposite directions [23, 27]. When these interactions are strong, the local pinning force *F*_pin_(*x*) can exceed the motor pulling force *F*_mot_, trapping the cargo in the off-state. In this state, the cargo undergoes diffusion-like motion near the microtubules for a duration *t*_off_, as illustrated in Figs. 4(a).

**FIG. 4.**
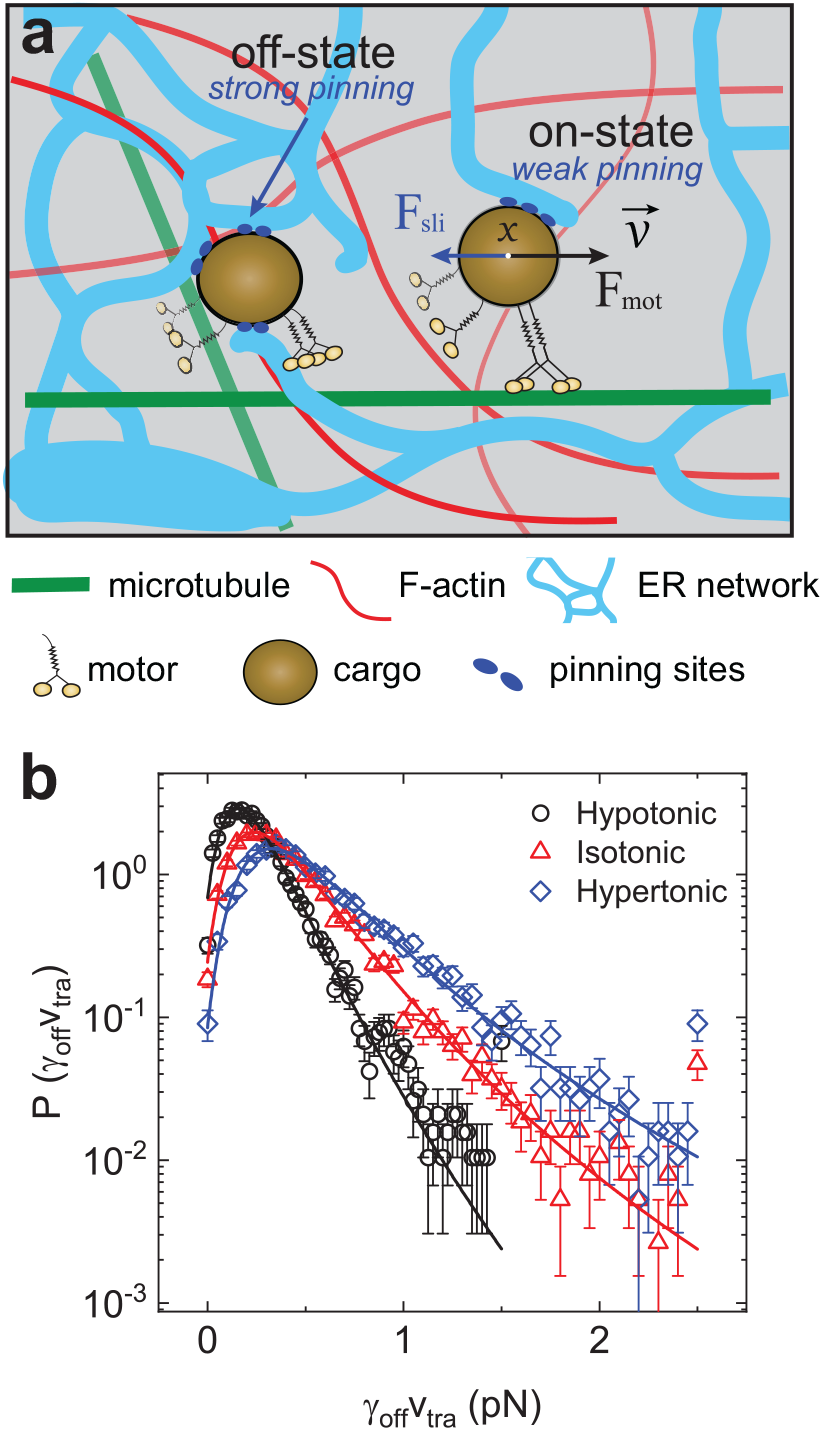
Stick-slip motion for cargo transport in living cells. (a) Schematic illustrating the two-state motion of cargo in a cell. (i) In the off-state (left), cargo is trapped by a strong pinning force *F*_pin_ from interactions with microtubule intersections (green lines), actin filaments (red lines), and the ER network (light blue). (ii) In the on-state (right), when the motor pulling force *F*_mot_ exceeds *F*_pin_, cargo undergoes continuous stick-slip motion, with a weaker sliding friction *F*_sli_. (b) Measured PDFs *P*(*γ*_off_*v*_tra_) for EGFR-endosomes in BEAS-2B cells treated with different osmotic solutions. The solid colored lines show the best fits of Eq. (5) to the data with the fitting parameters listed in Table S3.

A transition from static to kinetic friction occurs when the cargo finds a permissive path for directed motion along a microtubule. This transition is facilitated by non-equilibrium (ATP-dependent) network fluctuations [49] and remodeling [50, 51], or by enhanced motor cooperation [27, 52]. In the on-state, the cargo undergoes a continuous stick-slip motion, which can be described by the force balance equation:

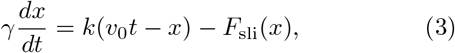

where *x*(*t*) represents the center of mass position of the cargo, which is pulled by an effective motor moving at a constant speed *v*_0_ along a microtubule, as illustrated in Fig. 4(a). The pulling force is transmitted to the cargo via a spring with spring constant *k*, and *v*_0_*t* represents the position of the effective motor. The symbol *F*_sli_(*x*) represents the sliding friction acting on the moving cargo at position *x*.

Using Eq. (3), we can make two predictions that can be directly tested in experiments. First, by averaging Eq. (3) across multiple mobile segments within each trajectory, we obtain the trajectory-based balance equation:

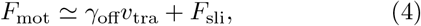

where *γ*_off_ = (*k*_*B*_*T*)*/D*_off_ is the measured drag coefficient from the off-state, with *k*_*B*_*T* being the thermal energy. Equation (4) indicates that the motor pulling force is dynamically balanced by two resistive forces. As the cargo moves, it must continuously break the pinning bonds with the surrounding protein networks, a process often referred to as pinning-depinning or stick-slip. Since the moving cargo has less time to interact with the surrounding protein networks, we have *F*_sli_ *< F*_pin_, as illustrated in Fig. 4(a). This situation resembles solid friction [53], where a mesoscale contact area slides over a rough surface, exhibiting stick-slip-like motion. In this case, the maximum forces needed to trigger local slips were found to follow a generalized extreme value (GEV) distribution [53, 54]. Therefore, we expect the viscous drag *γ*_off_*v*_tra_ also adheres to extreme value statistics.

To test this prediction, we calculate the trajectory-based diffusion coefficient (*D*_off_)_*j*_ in the off-state and convert it to the trajectory-based drag coefficient (*γ*_off_)_*j*_ = (*k*_*B*_*T*)*/*(*D*_off_)_*j*_ using the cell culture temperature *T* = 310 K (37^*°*^C) (see SI Sec. II.B. for more details [40]). With the trajectory-based *γ*_off_ and *v*_tra_, we compute the PDFs *P*(*γ*_off_*v*_tra_) for EGFR-endosomes in BEAS-2B cells treated with different osmotic solutions, as shown in Fig. 4(b). All three datasets exhibit an asymmetrical distribution with a long tail. Unlike the velocity PDFs, which have an exponential tail, the measured PDFs *P*(*γ*_off_*v*_tra_) decay more slowly than exponential and can be described by the GEV distribution [55],

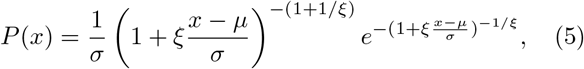

where *x* = *γ*_off_*v*_tra_, and *μ, σ*, and *ξ* are fitting parameters. The location parameter *μ* and scale parameter *σ* approximate the mean and standard deviation of the random variable *x*. This equation is commonly used to model the distribution of extreme values in independent and identically distributed random variables (see SI Sec. III.B for more details [40]). As shown in Fig. 4(b), the GEV distribution (solid lines) fits the data well, with the fitting parameters listed in Table S3.

The peak value of the measured *P*(*γ*_off_*v*_tra_) in the hypotonic solution is approximately 0.09 pN, increasing to 0.16 and 0.21 pN, as cell volume decreases. This result suggests that higher intracellular viscosity and viscoelasticity lead to greater drag forces, as expected. A previous study [49] showed that ATP depletion caused the diffusion coefficient of small inert particles in living cells to decrease 42-69 times, suggesting that the effective temperature *T* in living cells is elevated from the cell culture temperature by this factor, due to ATP-dependent active agitation. Multiplying the peak value of *γ*_off_*v*_tra_ by 42-69, we estimate that the actual peak value in the isotonic solution falls within the range of 6.7–11 pN. Assuming each motor protein exerts a force of 4 pN to pull the cargo [2], we estimate that 2-3 motor proteins contribute to the peak value of *γ*_off_*v*_tra_ in the isotonic solution. This estimate aligns with reported numbers of 1-10 motor proteins associated with each vesicle in cells [23, 56].

Our second prediction concerns the instantaneous cargo speed *v*(*t*), which can be derived directly from Eq. (3). Since the sliding frictional force *F*_sli_(*x*) varies randomly within a cell, the resulting cargo speed *v*(*t*) is expected to be a random variable. Therefore, we need to calculate the velocity PDF *P*(*v*). Because *F*_sli_(*x*) represents a cumulative sum of individual pinning forces on the cargo surface, it is spatially correlated across a range of cargo sizes. This correlation can be expressed mathematically as [57], ⟨|*F*_sli_(*x*) − *F*_sli_(*x*^*I*^)|^2^⟩ = 2*D*_F_|*x* − *x*^*I*^|, where *D*_F_ measures the friction roughness of *F*_sli_(*x*). This Brownian correlation is a general feature of the random pinning force field *F*_sli_(*x*) and is independent of its specific details [53]. Equation (3) has been used to model various physical problems related to random fields and is known as the Alessandro-Beatrice-Bertotti-Montorsi (ABBM) model [58–60].

We perform numerical simulations of Eq. (3) and confirm that the steady-state PDF *P*(*v*) follows a Gamma distribution, as described in Eq. (2). The Gamma distribution for *P*(*v*) can also be derived analytically by solving the Fokker-Planck equation related to Eq. (3) [59, 60], with *α* = 1*/D*^*I*^ and 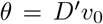, where 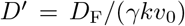 serves as the dimensionless control parameter in Eq. (3) (see SI Sec. III.A for more details [40]). When *α >* 1 (or *D*^*I*^ *<* 1), the Gamma distribution *P*(*v*) is a single peaked function for sliding stick-slip motion. At *α* = 1 (or *D*^*I*^ = 1), it simplifies to an exponential function. From the fitted values of *α* = 2.8 and *θ* = 0.35 presented in Fig. 2(c) (see Table S2 in SI), we find the motor pulling speed *v*_0_ = *αθ* = 0.98*v*_tra_. This result is expected, which further confirms our model.

The excellent agreement between the theoretical predictions of Eq. (3) and the fitting results in Figs. 2–4 (and in Figs. S2-S5 in SI [40]) demonstrates that the stick-slip model effectively captures the essential physics of cargo transport within cells. This model outlines a general mechanism for motor proteins to overcome various resistive forces and initiate directed cargo motion, following a univeral Gamma distribution. Its implications extend beyond cargo movement; it is vital for a range of cellular processes that reply on the intricate interplay between active (ATP-dependent) pulling and various resistances from the surrounding environment. Notable examples include vesicle budding [61], organelle positioning and fission [20], ER remodeling [62], cell migration [63, 64], and cancer cell invasion [65]. By quantitatively understanding these processes, the stick-slip model will provide valuable insights into the mechanics of cellular function and the underlying principles that govern life at the subcellular and cellular levels.

## Methods

Details about the materials and methods are given in Supplementary Materials.

## Supporting information

Supplemental Information concerning the article

## Data Availability

Source data for the figures in the main text and supplementary information are provided in this paper. Raw data generated in this study are available from the corresponding author (P.T.) upon request.

## Acknowledgments

The authors wish to thank Dr. Qirui Zhao for preparing the plasmids for cell transfections. This work was supported, in part, by RGC and ITC of Hong Kong SAR under grant nos. 16300224 (P.T.), C6041-24G-A (P.T.), 16104223 (P.H.), and ITCPD/17-9 (P.H.); by NSTC of Taiwan under grant no. 113-2112-M008-018-MY2 (P.Y.L.); and by NIH of USA under grant no. 2R35GM133688 (K.M. O.-M.).

## Author Contributions

Y.S. and P.T. designed the experiments; Y.S. performed the experiments; P.Y.L. and C.Y. developed the theoretical model with inputs from Y.S. and P.T.; all authors analyzed and interpreted data; Y.S. and P.T. drafted the paper with inputs from all authors; P.T. coordinated the project.

