## Supplemental Information concerning the article for "Stick-slip motion and universal statistics of cargo transport within living cells"

### Supplemental Information concerning the article: Stick-slip motion and universal statistics of cargo transport within living cells

Yusheng Shen,[1,3] Caishan Yan,[1] Pingbo Huang,[2] Kassandra M. Ori-McKenney,[3] Pik-Yin Lai,[4,5] Penger Tong[1]
*[1] Department of Physics and [2] Division of Life Science,*
*Hong Kong University of Science and Technology, Clear Water Bay, Kowloon, Hong Kong*
*[3] Department of Molecular and Cellular Biology,*
*University of California, Davis, Davis, CA 95616, USA*
*[4] Department of Physics and Center for Complex Systems,*
*National Central University, Taoyuan City 320, Taiwan*
*[5] Physics Division, National Center for Theoretical Sciences, Taipei 10617, Taiwan*
(Dated: March 20, 2025)

#### I. MATERIALS AND METHODS

**Cell culture, transfection and osmotic treatments.** BEAS-2B cells and HeLa cells, were cultured in Dulbecco's Modified Eagle Medium (DMEM, Life Technologies), and RPE-1 cells were cultured in Dulbecco's Modified Eagle Medium/Nutrient Mixture F-12 (DMEM/F-12, Life Technologies). Both media were supplemented with 10% fetal bovine serum (FBS), 50 units/mL of penicillin and 50 µg/mL of streptomycin. All cell cultures were maintained in a 95% air/5% $CO_2$ atmosphere at 37 °C. The cell lines were routinely confirmed to test negative for mycoplasma contamination. For live cell imaging, cells were seeded at a density of approximately $1 \times 10^4$ $cm^{-2}$ on a glass coverslip which was placed in a 35-mm polystyrene tissue-culture dish.

The expression vectors used in the study were EGFP-Rab5 (Addgene plasmid No. 49888; for expression of early-endosome marker, Rab5), LAMP1-mGFP (Addgene plasmid No. 34831, for expression of lysosome marker, LAMP1), and EGFP-Rab6A (Addgene plasmid No. 49469, for expression of secretory vesicle marker, Rab6A). Transfections were performed by using the Lipofectamine 2000 reagent kit (Invitrogen) according to manufacturer's instructions. Cells were generally transfected for 8 hrs with 1 $\mu$g plasmids when the density reached ∼80% confluency.

To manipulate the cytoplasm crowdedness, cells were treated with and imaged in extracellular osmotic environments ranging from hypoosmotic to hyperosmotic conditions (250–400 mOsm). These solutions were prepared by adding, respectively, 0.28 (∼400 mOsm, hypertonic), 0.2 (∼310 mOsm, isotonic control) and 0.15 M (∼250 mOsm, hypotonic) D-mannitol (Sigma) to a hypotonic base solution (in millimolar: 40 NaCl, 5 KCl, 1 $CaCl_2$, 2 $MgCl_2$, 10 HEPES (pH 7.4), ∼91 mOsm) to maintain a constant ionic strength. Imaging typically starts after 10 min of osmotic treatments.

**EGFR labelling.** The stock EGF-QD complex was prepared by mixing biotin-EGF (4 nM, Thermo Fisher Scientific) with streptavidin-conjungated quantum dots (QDs) with maximum fluorescent emission at 655 nm (QD655, 2 nM, Thermo Fisher Scientific) *in vitro* in culture medium at 4°C for 3 hrs. For high speed, short duration recording of EGFR-endosomes, EGFRs in the plasma membrane were labelled with EGF-QDs at a concentration of 0.8 nM biotin-EGF at room temperature for 5 min. After washing with culture medium for three times (5 min each), the cell-containing glass coverslip was then mounted on a coverslip holder (SC15012, Aireka Cells), which was finally mounted on an inverted microscope (DM-IRB, Leica) with a 100× objective (NA = 1.40). A live cell imaging chamber (CU-501, Chamlide TC) was equipped to the microscope to provide optimal culture conditions (95% air/5% $CO_2$ atmosphere at 37°C) for cells during imaging.

**Optical imaging and single-particle tracking.** EGFR-endosome, lysosome and early endosome were imaged using an inverted microscope (DM-IRB, Leica) with a 100× objective (NA = 1.40). Image sequences were typically recorded at 10 fps (frame per second) for 3 min by using an electron-multiplying charged-coupled device (EMCCD) camera (Ixon3 897, Andor). The exposure time for each frame is 30 ms and a microscope shutter was used to control the overall UV exposure time and reduce the light-induced damage to the living cells. The recorded images have 16 bits of gray scales and a spatial resolution of 512×512 pixels with the width of each pixel $p_w = 133$ nm in our optical setup.

Secretory vesicles were imaged using total internal reflection fluorescence (TIRF) microscopy, which was performed on an inverted research microscope Eclipse Ti2-E with the Perfect Focus System (Nikon), equipped with a 1.49 NA 100× TIRF objective with the 1.5× tube lens setting, a Ti-S-E motorized stage, piezo Z-control (Physik Instrumente), LU-N4 laser units (Nikon) as the light source, an iXon DU897 cooled EMCCD camera (Andor) with an high-speed emission filter wheel (ET480/40M for mTurquoise2, ET525/50M for GFP, ET520/40M for YFP, and ET632/60M for mRuby2; Chroma). The microscope was controlled with NIS Elements software (Nikon). Live cell imaging was performed in a live cell imaging chamber (H301-Nikon-TI-S-ER, Oko Labs) that was equipped to the microscope to provide optimal culture conditions (95% air/5% $CO_2$ atmosphere at 37 °C) for cells during imaging. The recorded images have 16

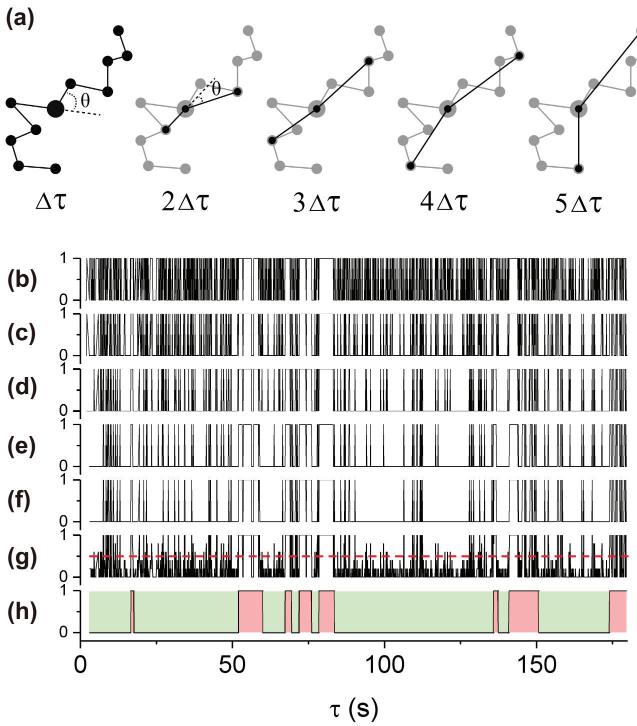


FIG. S1. **Sorting of the on-state and off-state segments.** (a) A schematic showing how to extract the turning angles, $\theta$, for a given point in a trajectory over 5 time steps. The point of concern and the angles are highlighted by a large-sized spot and two black lines, respectively. (b-f) Time series data of the discretized turning angles $\theta$ for a representative trajectory over 5 different time steps (from top to bottom: 0.1 s, 0.2 s, 0.3 s, 0.4 s, and 0.5 s). The five turning angles ranging from 0 (straight forward motion) to $\pi$ (straight backward motion) were discretized into two values: 1 if $\theta < \pi/2$ and 0 if $\theta \geq \pi/2$, without considering the left and right turns. (g) Time series data of the averaged value of the five discretized angles from (b-f). A threshold value of 0.5 (red dashed line) is used to partition the trajectory into the on-state segments (with $\langle\theta\rangle \geq 0.5$) and the off-state segments (with $\langle\theta\rangle < 0.5$). (h) Finally, the extracted on-state segments (red) and off-state segments (green) states after applying a sliding window to filter out some transient hopping events between the on-state and off-state.

bits of gray scales and a spatial resolution of 512×512 pixels with the width of each pixel $p_w = 107$ nm in our optical setup.

Single-particle tracking (SPT) was performed using a homemade tracking program written in MATLAB as previously described [1], which is based on the standard tracking algorithm [2, 3]. With this advanced SPT algorithm, we were able to obtain the position $\mathbf{r}(t)$ at time $t$ for EGFR-endosomes, early endosomes, lysosomes and secretory vesicles, and their trajectories were constructed from the consecutive images.

**Sorting of on-state and off-state segments.** To separate the on-state and off-state segments from each individual trajectory, we used an automated selection algorithm developed by Röding *et al.* [4] to evaluate the directional persistence of local movements, as illustrated in Fig. S1. Specifically, we analyzed the turning angles, $\theta$, around an arbitrary point in a given trajectory over a number of time steps (five time steps were used in this study) to determine the local directional persistence of the point. The five turning angles were discretized into two values: 1 if $\theta < \pi/2$ (considered as a forward movement without a turn) and 0 if $\theta \geq \pi/2$ (considered as a backward movement with a turn). The five discretized angles around the given point were then averaged and a threshold of 0.5 was used to determine whether the segment of the trajectory around the given point belongs to the on-state (with $\langle\theta\rangle \geq 0.5$) or the off-state (with $\langle\theta\rangle < 0.5$). Finally, a sliding window of 10 time steps (1 s) was applied to the discretized time series to filter out some transient hopping events between the on-state and off-state to finalize the on-state and off-state segments. In the experiment, only those on-state segments with a duration longer than 0.3 s were considered as the true on-state segments.

**Summary of sample sizes.** Details about the sample sizes used in this study are summarized in Table S1. To build up the statistics, we typically track each type of vesicles from more than 35 cells cultured under the same condition so that more than $4\times10^3$ trajectories were used for statistical analysis.

#### II. SUPPLEMENTARY FIGURES AND TABLES

##### A. Statistics of cargo velocities

In addition to the velocity measurements for EGFR-endosomes in live BEAS-2B cells presented in Fig. 2 of the main text, we also investigate the PDFs of the on-state velocities, including $v_{\rm seg}$, $v_{\rm seg}/v_{\rm tra}$, and $v_{\rm ins}/v_{\rm seg}$, for other vesicles across various cell types and levels of intracellular crowdedness. Figure S2 shows the PDFs $P(v_{\rm seg})$ of segment velocity $v_{\rm seg}$ for EGFR-endosomes in live BEAS-2B cells treated with different osmotic solutions (Fig. S2(a)), across various cell types (Fig. S2(b)), and for different vesicles in live BEAS-2B cells (Fig. S2(c)). Consistent with our findings for EGFR-endosomes in BEAS-2B cells, the measured segment velocity $v_{\rm seg}$ shows substantial variations across different conditions, and its PDFs $P(v_{\rm seg})$ can be well described by the Gamma distribution (solid lines in Fig. S2), with the fitting parameters detailed in Table S2.

Furthermore, when the velocity fluctuations among different trajectories are normalized by their mean value $v_{\rm tra}$, the resulting PDFs $P(v_{\rm seg}/v_{\rm tra})$ converge to a common Gaussian distribution curve (red solid line in Fig. S3(a)). The fitted mean and standard deviation for vesicles under various conditions are given in Table S2. It is found that all the PDFs $P(v_{\rm seg}/v_{\rm tra})$ have a similar

| Cell type | Vesicle type | Cell treatment | No. of cells | No. of mobile trajectories |
|---|---|---|---|---|
| BEAS-2B | EGFR-endosome | No treatment | 83 | 9946 |
| BEAS-2B | Lysosome | No treatment | 42 | 17465 |
| BEAS-2B | Early endosome | No treatment | 39 | 20780 |
| BEAS-2B | Secretory vesicle | No treatment | 35 | 19106 |
| HeLa | EGFR-endosome | No treatment | 51 | 6007 |
| RPE-1 | EGFR-endosome | No treatment | 44 | 7917 |
| BEAS-2B | EGFR-endosome | Hypotonic | 63 | 9217 |
| BEAS-2B | EGFR-endosome | Isotonic | 49 | 8846 |
| BEAS-2B | EGFR-endosome | Hypertonic | 48 | 4752 |
| | | Total: | 454 | 104036 |

TABLE S1. Summary of cell treatments and sample sizes. The columns, from left to right, provide information on the cell types used, the types of vesicles imaged, the treatments applied to the cells, the number of cells that were imaged, and the total number of mobile trajectories included in the statistical analysis.

| *Vesicle type, cell type & treatment* | | EGFR-endosome BEAS-2B | Lysosome BEAS-2B | Early endosome BEAS-2B | Secretory vesicle BEAS-2B | EGFR-endosome HeLa | EGFR-endosome RPE-1 | EGFR-endosome BEAS-2B hypotonic | EGFR-endosome BEAS-2B isotonic | EGFR-endosome BEAS-2B hypertonic |
|---|---|---|---|---|---|---|---|---|---|---|
| $v_{tra}$ (μm/s) | $\alpha$ | 4.7 ± 0.05 | 4.7 ± 0.06 | 3.1 ± 0.05 | 5.9 ± 0.06 | 4.7 ± 0.05 | 3.6 ± 0.04 | 3.6 ± 0.06 | 3.2 ± 0.05 | 2.8 ± 0.06 |
| | $\theta$ | 0.18 ± 0.01 | 0.21 ± 0.02 | 0.31 ± 0.02 | 0.11 ± 0.02 | 0.18 ± 0.01 | 0.25 ± 0.01 | 0.25 ± 0.02 | 0.22 ± 0.01 | 0.20 ± 0.02 |
| | $\alpha\theta$ | 0.85 ± 0.05 | 0.99 ± 0.09 | 0.96 ± 0.06 | 0.65 ± 0.12 | 0.85 ± 0.05 | 0.90 ± 0.04 | 0.90 ± 0.07 | 0.70 ± 0.03 | 0.56 ± 0.06 |
| | $\sqrt{\alpha}\theta$ | 0.39 ± 0.02 | 0.46 ± 0.04 | 0.55 ± 0.04 | 0.27 ± 0.05 | 0.39 ± 0.02 | 0.47 ± 0.02 | 0.47 ± 0.04 | 0.39 ± 0.02 | 0.33 ± 0.03 |
| $v_{seg}$ (μm/s) | $\alpha$ | 2.95 ± 0.05 | 3.4 ± 0.06 | 2.4 ± 0.06 | 4.4 ± 0.05 | 3.9 ± 0.06 | 2.8 ± 0.06 | 2.7 ± 0.06 | 2.2 ± 0.06 | 1.9 ± 0.08 |
| | $\theta$ | 0.28 ± 0.01 | 0.29 ± 0.02 | 0.38 ± 0.02 | 0.17 ± 0.01 | 0.21 ± 0.01 | 0.30 ± 0.01 | 0.33 ± 0.02 | 0.29 ± 0.02 | 0.27 ± 0.02 |
| | $\alpha\theta$ | 0.83 ± 0.03 | 0.99 ± 0.07 | 0.91 ± 0.05 | 0.75 ± 0.04 | 0.82 ± 0.04 | 0.84 ± 0.03 | 0.89 ± 0.06 | 0.64 ± 0.05 | 0.51 ± 0.04 |
| | $\sqrt{\alpha}\theta$ | 0.48 ± 0.02 | 0.53 ± 0.04 | 0.59 ± 0.03 | 0.36 ± 0.02 | 0.41 ± 0.02 | 0.50 ± 0.02 | 0.54 ± 0.03 | 0.43 ± 0.03 | 0.37 ± 0.03 |
| $v_{ins}/v_{seg}$ | $\alpha$ | 2.8 ± 0.05 | 2.9 ± 0.06 | 2.8 ± 0.05 | 2.4 ± 0.08 | 3.4 ± 0.08 | 3.6 ± 0.06 | 3.1 ± 0.05 | 2.8 ± 0.08 | 2.7 ± 0.06 |
| | $\theta$ | 0.35 ± 0.01 | 0.36 ± 0.02 | 0.37 ± 0.01 | 0.43 ± 0.03 | 0.30 ± 0.03 | 0.29 ± 0.01 | 0.33 ± 0.02 | 0.36 ± 0.03 | 0.38 ± 0.02 |
| | $\alpha\theta$ | 0.98 ± 0.03 | 1.04 ± 0.06 | 1.04 ± 0.03 | 1.03 ± 0.08 | 1.02 ± 0.10 | 1.04 ± 0.04 | 1.02 ± 0.06 | 1.01 ± 0.09 | 1.03 ± 0.06 |
| | $\sqrt{\alpha}\theta$ | 0.59 ± 0.02 | 0.61 ± 0.03 | 0.62 ± 0.02 | 0.67 ± 0.05 | 0.55 ± 0.06 | 0.55 ± 0.02 | 0.58 ± 0.04 | 0.60 ± 0.05 | 0.62 ± 0.03 |
| $v_{seg}/v_{tra}$ | $\mu$ | 0.98 ± 0.02 | 0.96 ± 0.03 | 0.94 ± 0.03 | 0.98 ± 0.02 | 0.97 ± 0.02 | 0.96 ± 0.03 | 0.95 ± 0.03 | 0.93 ± 0.02 | 0.94 ± 0.02 |
| | $\sigma$ | 0.64 ± 0.01 | 0.70 ± 0.02 | 0.69 ± 0.03 | 0.65 ± 0.01 | 0.63 ± 0.01 | 0.60 ± 0.01 | 0.69 ± 0.02 | 0.69 ± 0.03 | 0.64 ± 0.02 |

TABLE S2. **Summary of fitting results for cargo velocities.** The first three rows present the best-fit values of the parameters $\alpha$ and $\theta$ for the Gamma distribution described in Eq. (2) of the main text, applied to the measured velocity PDFs of various vesicle types, cell types, and treatments. For reference, we also include the mean ($\alpha\theta$) and standard deviation ($\sqrt{\alpha}\theta$) of the fitted cargo velocities. The fourth row provides the best-fit values of the parameters $\mu$ (mean value) and $\sigma$ (standard deviation) for the Gaussian fit, $P(v) \propto \exp[-(v-\mu)^2/(2\sigma^2)]$, applied to the measured velocity PDFs across different vesicle types, cell types, and treatments. Error bars indicate the fitting uncertainties.

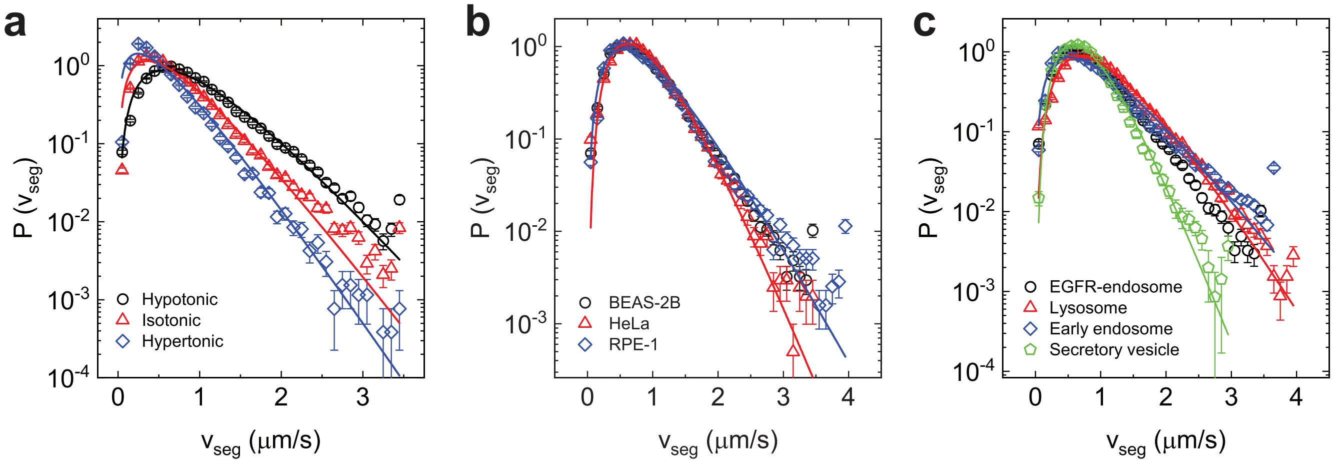


FIG. S2. **Statistics of on-state segment velocity $v_{\rm seg}$.** (a) Measured PDFs $P(v_{\rm seg})$ for EGFR-endosomes in live BEAS-2B cells treated with hypotonic (black circles), isotonic (red triangles), and hypertonic (blue diamonds) solutions. (b) Measured PDFs $P(v_{\rm seg})$ for EGFR-endosomes in live BEAS-2B cells (black circles, re-plot from Fig. 2(a) of the main text for comparison), HeLa cells (red triangles) and RPE-1 cells (blue diamonds). (c) Measured PDFs $P(v_{\rm seg})$ for EGFR-endosomes (black circles, re-plot from Fig. 2(a) of the main text for comparison), lysosomes (red triangles), early endosomes (blue diamonds) and secretory vesicles (green pentagons) in live BEAS-2B cells. The colored solid lines in panels (a), (b), and (c) show the best fits of the Gamma distribution described in Eq. (2) of the main text to the data points, with the fitting parameters listed in Table S2.

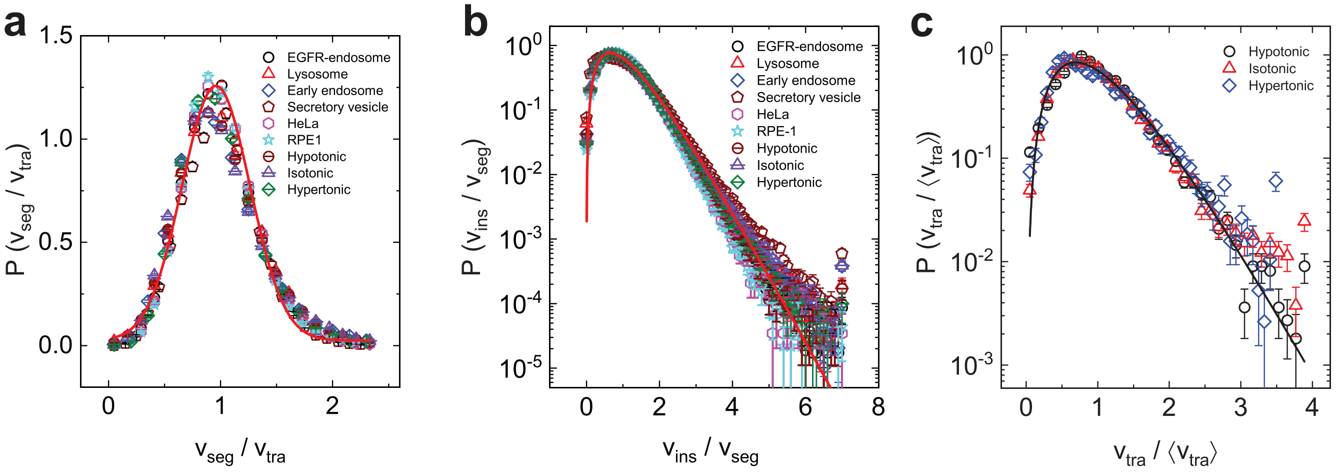


FIG. S3. **Universal statistics of on-state velocities for different vesicles, cell types, and intracellular crowdedness.** (a) Measured PDFs $P(v_{\rm seg}/v_{\rm tra})$ of the normalized segment velocity, $v_{\rm seg}/v_{\rm tra}$. For simplicity, only the Gaussian fit, $P(v) \propto \exp[-(v-\mu)^2/(2\sigma^2)]$, is shown for the data points corresponding to EGFR-endosomes in HeLa cells (red solid line). The mean value $\mu$ and standard deviation $\sigma$ are listed in Table S2. (b) Measured PDFs $P(v_{\rm ins}/v_{\rm seg})$ of the normalized instantaneous velocity, $v_{\rm ins}/v_{\rm seg}$. For simplicity, only the fit of the Gamma function described in Eq. (2) of the main text is shown for the data points corresponding to EGFR-endosomes in BEAS-2B cells treated with isotonic solutions (red solid line). The fitting parameters are provided in Table S2. The measurements in panels (a) and (b) include EGFR-endosomes, lysosomes, early endosomes, and secretory vesicles in live BEAS-2B cells, as well as EGFR-endosomes in live HeLa and RPE-1 cells, and EGFR-endosomes in live BEAS-2B cells treated with different osmotic solutions. (c) Measured PDFs $P(v_{\rm tra}/\langle v_{\rm tra}\rangle)$ of the normalized trajectory velocity, $v_{\rm tra}/\langle v_{\rm tra}\rangle$, for EGFR-endosomes in live BEAS-2B cells treated with various osmotic solutions. The black solid line shows the best fit of Eq. (2) of the main text to the data points, with fitting parameters $\alpha = 3.28 \pm 0.01$ and $\theta = 0.30 \pm 0.01$.

mean value close to unity ($\mu \simeq 1$) and a standard deviation of about $\sigma \simeq 0.65$. This result strongly supports the notion that, for a given trajectory (or motor-cargo complex), the intermittent switching between the on-state and off-state does not significantly alter the vesicle velocity across different segments. This characteristic appears to be universal across various vesicles, cell types, and intracellular environments. The relatively stable segment velocity $v_{\rm seg}$ within individual trajectories indicates that the motor-cargo complexes remain stable during long-

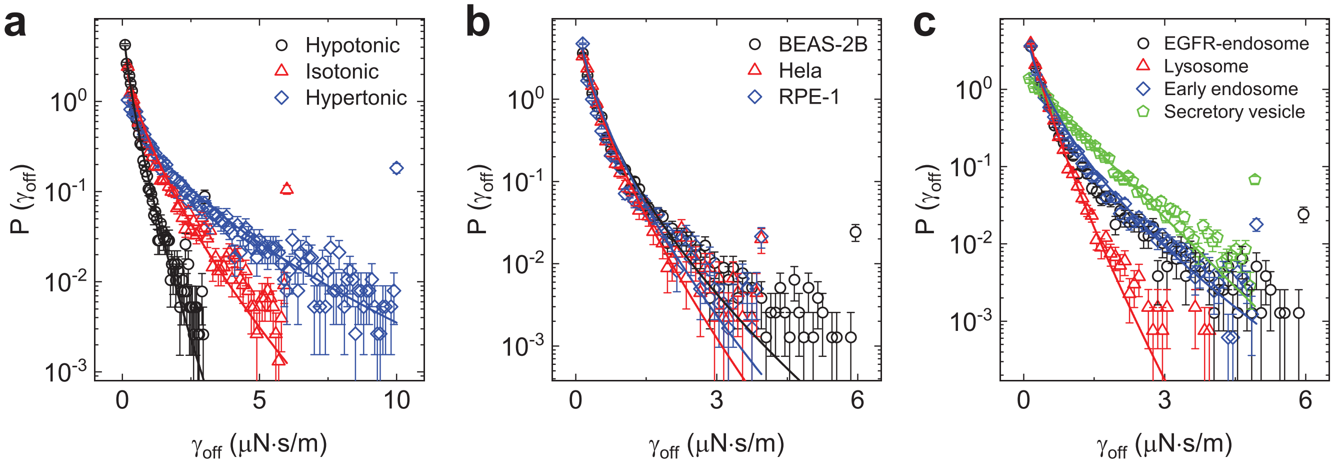


FIG. S4. **Statistics of drag coefficient $\gamma_{off}$.** (a) Measured PDFs $P(\gamma_{off})$ for EGFR-endosomes in live BEAS-2B cells treated with hypotonic (black circles), isotonic (red triangles), and hypertonic (blue diamonds) solutions. (b) Measured PDFs $P(\gamma_{off})$ for EGFR-endosomes in live BEAS-2B cells (black circles), HeLa cells (red triangles), and RPE-1 cells (blue diamonds). (c) Measured PDFs $P(\gamma_{off})$ for EGFR-endosomes (black circles), lysosomes (red triangles), early endosomes (blue diamonds), and secretory vesicles (green pentagons) in live BEAS-2B cells. The colored solid lines in (a), (b), and (c) show the stretched exponential fits, $P(\gamma_{off}) \propto \exp[-(\gamma_{off}/\gamma_0)^\beta]$, to the data points, with the fitting parameters provided in Table S3.

distance transport, even in the presence of a heterogeneous cellular environment.

Figure S3(b) shows the measured PDFs $P(v_{ins}/v_{seg})$ for various vesicles across different cell types and levels of intracellular crowdedness. All the PDFs $P(v_{ins}/v_{seg})$ converge to a common Gamma distribution curve (red solid line). From the fitted values of $\alpha$ and $\theta$, we find the motor pulling speed $v_0 = \alpha\theta$ (see Eq. (S7) below) falls within a narrow range of 0.98 to 1.04 times $v_{tra}$. These results further confirms that the stick-slip model effectively captures the essential physics of cargo transport in living cells.

Figure S3(c) shows the measured PDFs $P(v_{tra}/\langle v_{tra}\rangle)$ of the normalized trajectory velocity, $v_{tra}/\langle v_{tra}\rangle$, for EGFR-endosomes in BEAS-2B cells treated with different osmotic solutions. The PDFs $P(v_{tra}/\langle v_{tra}\rangle)$ collapse onto a common Gamma distribution curve (black solid line) when the velocity variable is normalized by its mean value $\langle v_{tra}\rangle$. Figure S3(c) thus suggests that the observed Gamma distribution of $v_{tra}$ is a robust feature in live BEAS-2B cells, independent of intracellular crowdedness.

Table S2 summarizes the fitting results presented in Figs. 2–4 of the main text, as well as in Figs. S2 and S3. Analysing the fitting parameters in Table S2 reveals several interesting features. First, the measured $v_{seg}$ and $v_{tra}$ show a comparable standard deviation, suggesting that fluctuations of $v_{seg}$ result primarily from dynamic heterogeneity in $v_{tra}$. Second, secretory vesicles, which are primarily transported by kinesin motors [5], exhibit a significantly lower mean velocity $\alpha\theta$ than other vesicle types, whose transport also involves dynein motors. Additionally, secretory vesicles have a smaller standard deviation $\sqrt{\alpha}\theta$ compared to other vesicles. This finding is consistent with previous research indicating that kinesin-based vesicle movements are slower and more homogeneous than dynein-based vesicle movements [6]. Third, both the mean ($\alpha\theta$) and standard deviation ($\sqrt{\alpha}\theta$) of trajectory velocity $v_{tra}$ for EGFR-endosomes in BEAS-2B cells show a decrease as intracellular crowdedness increases. The observed changes in velocity statistics may be attributed to variations in the effective viscosity of the cytoplasm under different osmotic conditions.

##### B. Statistics of drag coefficient $\gamma_{off}$ and viscous drag force $\gamma_{off}v_{tra}$

To investigate the statistics of viscous drag $\gamma_{off}v_{tra}$, we first study the statical properties of drag coefficient $\gamma_{off}$. Figure S4 shows the measured PDFs $P(\gamma_{off})$ for EGFR-endosomes in live BEAS-2B cells treated with different osmotic solutions (Fig. S4(a)), across various cell types (Fig. S4(b)), and for different vesicles in live BEAS-2B cells (Fig. S4(c)). All the PDFs $P(\gamma_{off})$ exhibit a long decay tail and are well described by a stretched exponential distribution (solid lines):

$$P(\gamma_{off}) \propto e^{-(\gamma_{off}/\gamma_0)^\beta}, \tag{S1}$$

where $\gamma_0$ and $\beta$ are two fitting parameters. The best-fit values of $\gamma_0$ and $\beta$ for vesicles under various conditions are given in Table S3. The stretched exponential distribution is commonly used to characterize dynamic heterogeneity associated with individual trajectories, which vary significantly with different values of $p_{on}$ (see Fig. 1(c) in the main text). The variations in $\gamma_{off}$ may result from spatial variations in cytoplasm crowdedness [7], active agitations through ATP-dependent processes, and sample variations of motor-cargo complexes.

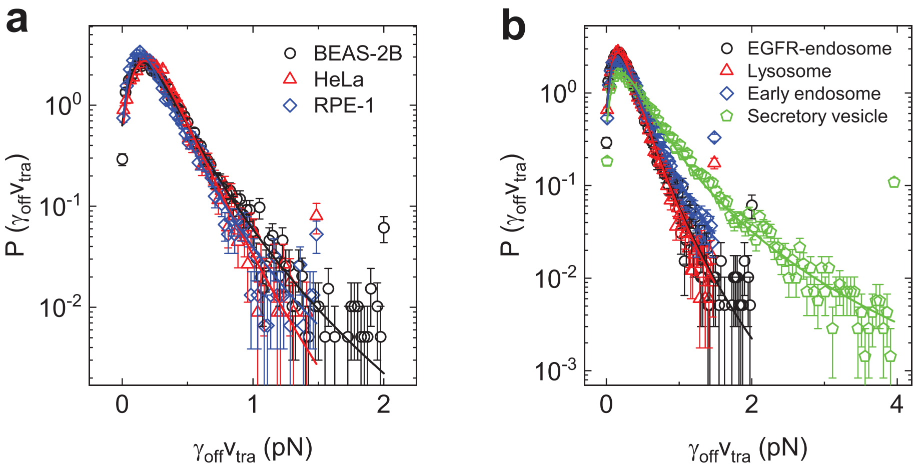


FIG. S5. **Statistics of viscous drag force $\gamma_{\mathrm{off}} v_{\mathrm{tra}}$.** (a) Measured PDFs $P(\gamma_{\mathrm{off}} v_{\mathrm{tra}})$ for EGFR-endosomes in live BEAS-2B cells (black circles), HeLa cells (red triangles), and RPE-1 cells (blue diamonds). (b) Measured PDFs $P(\gamma_{\mathrm{off}} v_{\mathrm{tra}})$ for EGFR-endosomes (black circles), lysosomes (red triangles), early endosomes (blue diamonds), and secretory vesicles (green pentagons) in live BEAS-2B cells. The solid colored lines in (a) and (b) show the best fits of the GEV distribution described in Eq. (4) of the main text to the data points, with the fitting parameters listed in Table S3.

| *Vesicle type, cell type & treatment* | | EGFR-endosome BEAS-2B | Lysosome BEAS-2B | Early endosome BEAS-2B | Secretory vesicle BEAS-2B | EGFR-endosome HeLa | EGFR-endosome RPE-1 | EGFR-endosome BEAS-2B hypotonic | EGFR-endosome BEAS-2B isotonic | EGFR-endosome BEAS-2B hypertonic |
|---|---|---|---|---|---|---|---|---|---|---|
| $\gamma_{\mathrm{off}}$ (μN·s/m) | $\gamma_0$ | 0.07 ± 0.01 | 0.12 ± 0.01 | 0.09 ± 0.01 | 0.61 ± 0.01 | 0.11 ± 0.01 | 0.09 ± 0.01 | 0.12 ± 0.01 | 0.22 ± 0.01 | 0.34 ± 0.01 |
| | $\beta$ | 0.56 ± 0.05 | 0.75 ± 0.04 | 0.56 ± 0.03 | 0.9 ± 0.05 | 0.67 ± 0.04 | 0.62 ± 0.04 | 0.70 ± 0.05 | 0.64 ± 0.04 | 0.55 ± 0.05 |
| $\gamma_{\mathrm{off}} v_{\mathrm{tra}}$ (pN) | $\xi$ | 0.17 ± 0.02 | 0.16 ± 0.02 | 0.29 ± 0.03 | 0.36 ± 0.02 | 0.07 ± 0.02 | 0.20 ± 0.02 | 0.09 ± 0.02 | 0.16 ± 0.02 | 0.21 ± 0.03 |
| | $\mu$ | 0.18 ± 0.01 | 0.18 ± 0.01 | 0.19 ± 0.01 | 0.27 ± 0.01 | 0.18 ± 0.01 | 0.15 ± 0.01 | 0.16 ± 0.01 | 0.27 ± 0.01 | 0.37 ± 0.01 |
| | $\sigma$ | 0.14 ± 0.01 | 0.14 ± 0.01 | 0.14 ± 0.01 | 0.24 ± 0.01 | 0.13 ± 0.01 | 0.12 ± 0.01 | 0.13 ± 0.01 | 0.19 ± 0.01 | 0.25 ± 0.01 |

TABLE S3. **Summary of fitting results for drag coefficient $\gamma_{\mathrm{off}}$ and viscous drag force $\gamma_{\mathrm{off}} v_{\mathrm{tra}}$.** The first row presents the best-fit values of the parameters $\gamma_0$ and $\beta$ for the stretched exponential distribution described in Eq. (S1), applied to the measured PDFs $P(\gamma_{\mathrm{off}})$ from various vesicle types, cell types, and treatments. The last row provides the best-fit values of the parameters $\xi$, $\mu$ and $\sigma$ for the GEV distribution described in Eq. (4) of the main text, applied to the measured PDFs $P(\gamma_{\mathrm{off}} v_{\mathrm{tra}})$ across different vesicle types, cell types, and treatments. Error bars indicate the fitting uncertainties.

Notably, the mean value of $\gamma_{\mathrm{off}}$, which correlates with cytoplasmic viscosity, is found to increase as cell volume decreases, as shown in Fig. S4(a). It is also found that $\gamma_{\mathrm{off}}$ varies widely for different vesicles in BEAS-2B cells (Fig. S4(c)), but remains relatively unchanged for the same vesicle in different cell types (Fig. S4(b)).

Figure S5 shows the measured PDFs $P(\gamma_{\mathrm{off}} v_{\mathrm{tra}})$ for EGFR-endosomes across different cell types (Fig. S5(a)) and for various vesicles in live BEAS-2B cells (Fig. S5(b)). All the PDFs $P(\gamma_{\mathrm{off}} v_{\mathrm{tra}})$ conform to extreme value statistics, as indicated by the solid lines. The best-fit values of $\xi$, $\mu$, and $\sigma$ for the vesicles under various conditions are provided in Table S3. The shape parameter $\xi$ influences the tail shape of the measured $P(\gamma_{\mathrm{off}} v_{\mathrm{tra}})$. When $\xi = 0$, the generalized extreme value (GEV) distribution in Eq. (4) of the main text is reduced to the Gumbel distribution (see Eq. (S9) below), which features a long exponential tail for large values of $\gamma_{\mathrm{off}} v_{\mathrm{tra}}$. Except for EGFR-endosomes in Hela cells and in BEAS-2B cells treated with a hypotonic solution – where the fitted value of $\xi$ is very small – the valued of $\xi$ for other vesicles across different conditions deviate from the $\xi = 0$ limit. In particular, we observe $\xi = 0.36$ for secretory vesicles in BEAS-2B cells.

As shown in Fig. 4(b) of the main text, the fitted value of $\xi$ gradually increases as cell volume decreases. By comparing Fig. S5(a) with Fig. S5(b), we find that the viscous drag forces for secretory vesicles, which are primarily transported by kinesins motors [5], are significantly higher than the rest endolysosomes (include EGFR-endosomes, lysosomes and early endosomes), whose transport also involves dynein motors. Collectively, Figs. S4 and S5 demonstrate that viscous drag force on vesicles adheres to extreme value statistics, strongly supporting the idea that the stick-slip model effectively captures the essential physics of cargo transport in living cells.

#### III. THEORETICAL ANALYSES

##### A. Stick-slip motion

We consider a phenomenological model to describe the vesicle motion, where an effective motor pulls the vesicle through a quasi-one-dimensional (1D) random pinning force field along a microtubule. In this model, we assume that the effective motor moves at a constant speed $v_0$, and the force is transmitted to the vesicle through a spring with a spring constant $k$. The equation of motion for the vesicle can be expressed as:

$$\gamma \frac{dx}{dt} = k(v_0 t - x) - F_{\text{sli}}(x), \tag{S2}$$

where $x$ represents the center-of-mass position of vesicle. On the left side of Eq. (S2), the term $\gamma(dx/dt)$ accounts for viscous damping from the cytoplasm, with $\gamma$ being the drag coefficient. On the right side, the first term $k(v_0t - x)$ represents the elastic pulling force exerted by the effective motor, with $v_0 t$ indicating the position of the motor. The second term, $F_{\text{sli}}(x)$ (which is greater than zero), represents the total pinning force against the motion, which results from various obstacles that interact with the moving vesicle.

In this model, we assume that vesicle motion is overdamped, so that the net driving force is always balanced by viscous damping and elastic friction. The pinning force $F_{\text{sli}}(x)$ has a positive mean value $F_0 = \langle F_{\text{sli}}(x)\rangle > 0$, which causes a constant mean stretch of $F_0/k$ in the pulling spring but does not affect the dynamics or velocities of the vesicle. The fluctuations in cargo velocity observed in the experiments are determined primarily by the fluctuating component $F_{\text{sli}}(x) - F_0$. Furthermore, we assume that $F_{\text{sli}}(x)$ exhibits Brownian correlation, characterized by the relation $\langle |F_{\text{sli}}(x) - F_{\text{sli}}(x')|^2\rangle = 2D_F|x - x'|$, where $D_F$ quantifies the amplitude of fluctuations in $F_{\text{sli}}(x)$. This Brownian correlation is considered a universal property of multi-site pinning force field [8] and has been utilized in modeling various random force fields. Equation (S2) was originally proposed to describe domain wall motion in magnetic systems [9] and is known as the Alessandro-Beatrice-Bertotti-Montorsi (ABBM) model [10, 11].

To obtain the governing equation of the velocity $v \equiv dx/dt$, we first differentiate both sides of Eq. (S2) with respect to $x$. Using $d/dx = \frac{1}{v}\frac{d}{dt}$, we have

$$\gamma \frac{dv}{vdt} = k(\frac{v_0}{v} - 1) + w(x), \tag{S3}$$

where $w(x) = dF_{\text{sli}}(x)/dx$ has a zero mean and obeys the relation $\langle w(x)w(x')\rangle = 2D_F\delta(x - x')$. Multiplying

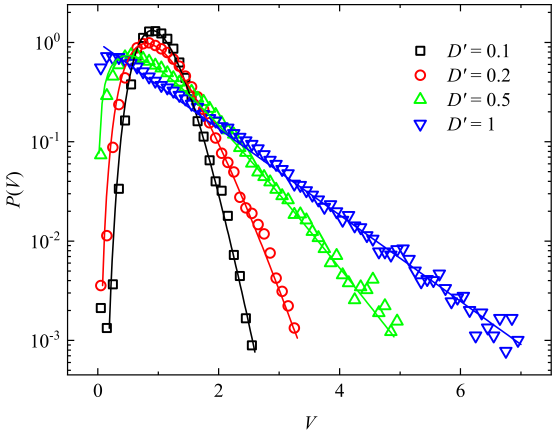


FIG. S6. **Numerical results.** Numerically calculated PDFs $P(V)$ of dimensionless velocity $V$ for different values of $D'$. The colored symbols are obtained from the Brownian dynamics simulations of Eq. (S5). The colored solid lines show the analytical PDFs shown in Eq. (S6) for different values of $D'$.

Eq. (S3) by $v$, we obtain

$$\gamma \frac{dv}{dt} = k(v_0 - v) + \sqrt{v}\eta(t), \tag{S4}$$

where $\eta(t) = \sqrt{v}w$ is a Gaussian white noise, having a zero mean and a correlation $\langle \eta(t)\eta(t')\rangle = 2D_F\delta(t - t')$.

Equation (S4) can be further simplified by making a variable replacement with $v = Vv_0$ and $t = T(\gamma/k)$. After the simplification, we obtain the dimensionless equation

$$\frac{dV}{dT} = 1 - V + \sqrt{V}\zeta(T), \tag{S5}$$

where $\langle \zeta(T)\zeta(T')\rangle = 2D'\delta(T - T')$ with $D' = D_F/(\gamma k v_0)$. Depending on the magnitude of the dimensionless control parameter $D'$, Eq. (S5) (or Eq. (S2)) yields two types of motion. When $D' < 1$, the motion is stable, in which $V$ vaires smoothly with time $T$. For $D' > 1$, the motion becomes unstable, in which velocity bursts ($V \gg 1$) occur intermittently over a background of slow motion with $V \ll 1$.

In both cases, the steady-state velocity distribution $P(V)$ can be obtained by solving the Fokker-Planck equation associated with Eq. (S5) and the final result is given by [9, 11]

$$P(V) = \frac{1}{D'^{1/D'}\Gamma(1/D')} V^{-1+1/D'} e^{-V/D'}. \tag{S6}$$

Equation (S6) can be rewritten in a dimensional form

$$p(v) = \frac{1}{\theta^\alpha \Gamma(\alpha)} v^{\alpha-1} e^{-v/\theta}, \tag{S7}$$

where $\alpha = 1/D'$, $\theta = D'v_0$, and $\Gamma(\alpha)$ is the Gamma function. Equation (S7) is provided in the main text as Eq. (2).

Equation (S6) reveals that $P(V)$ is a single peaked function for a stable sliding motion with $D' < 1$ (or $\alpha > 1$). When $D' > 1$ (or $\alpha < 1$), $P(V)$ decreases monotonically with $V$. In particular, when $D' = 1$ (or $\alpha = 1$), $P(V)$ follows the simple exponential distribution. Figure S6 shows the numerically calculated $P(V)$ from the Brownian dynamics simulations of Eq. (S5). The results in the regime of stable sliding motion agree well with the analytical prediction of Eq. (S6) (colored solid lines).

##### B. Extreme value statistics

The generalized extreme value (GEV) distribution [12] is presented in Eq. (5) of the main text and is reiterated here for convenience:

$$P(x) = \frac{1}{\sigma}\left(1+\xi\frac{x-\mu}{\sigma}\right)^{-(1+1/\xi)} e^{-(1+\xi\frac{x-\mu}{\sigma})^{-1/\xi}}, \quad \text{(S8)}$$

where $x$ $(= \gamma_{\rm off}v_{\rm tra})$ is a random variable. The GEV distribution is characterized by three independent parameters: a location parameter $\mu$, a scale parameter $\sigma$, and a shape parameter $\xi$. The location parameter $\mu$ and the scale parameter $\sigma$ are approximately the mean and standard deviation of $x$, respectively. The shape parameter $\xi$ determines the tail shape of the GEV distribution.

When $\xi > 0$, $P(x)$ exhibits a heavy tail that decays more slowly than an exponential tail, as shown in Fig. 4 of the main text, with the decay rate decreasing as $\xi$ increases. When $\xi = 0$, $P(x)$ has a long exponential tail. In this case, $P(x)$ takes on a much simpler form, which can be otained by taking the limit as $\xi \to 0$, i.e.,

$$\lim_{\xi\to 0} P(x) = \frac{1}{\sigma}\exp\left(-\frac{x-\mu}{\sigma} - e^{-(x-\mu)/\sigma}\right). \quad \text{(S9)}$$

Equation (S9) is also called the Gumbel distribution. When $\xi < 0$, $P(x)$ has a short tail or no tail.
